# A database of simulated tumor genomes towards accurate detection of somatic small variants in cancer

**DOI:** 10.1101/261503

**Authors:** Jing Meng, Yi-Ping Phoebe Chen

## Abstract

**Background:** Somatic mutations promote the transformation of normal cells to cancer. Accurate identification of such mutations facilitates cancer diagnosis and treatment. A number of callers have been developed to predict them from paired tumor/normal or unpaired tumor sequencing data. However, the small size of currently available experimentally validated somatic sites limits evaluation and then improvement of callers. Fortunately, NIST reference material NA12878 genome has been well-characterized with publicly available high-confidence genotype calls.

**Results:** We used BAMSurgeon to create simulated tumors by introducing somatic small variants (SNVs and small indels) into homozygous reference or wildtype sites of NA12878. We generated 135 simulated tumors from 5 pre-tumors/normals. These simulated tumors vary in sequencing and subsequent mapping error profiles, read length, the number of sub-clones, the VAF, the mutation frequency across the genome and the genomic context. Furthermore, these pure tumor/normal pairs can be mixed at desired ratios within each pair to simulate sample contamination.

**Conclusions:** This database (a total size of 15 terabytes) will be of great use to benchmark somatic small variant callers and guide their improvement.

## Background

Somatic mutations promote the transformation of normal cells to cancer [1-3]. Like germline mutations, the length of affected nucleotide sequences exclusively in cancer cells ranges from one nucleotide to entire chromosomes [4, 5]. The ultimate goal of cancer research is precise therapeutic targeting. To achieve the goal, a series of studies have been conducting, including but not limited to: identifying genes that drive cancer progression [6-8]; classification of cancer subtypes to establish the correlation between molecular properties and clinical outcomes [9, 10]; and linking environmental factors to mutational patterns in cancer genomes [11, 12]. Accurate identification of somatic mutations is the first step to therapeutic precision, which is followed by the aforementioned studies, and plays a key role in clinical diagnosis.

In an ideal error-free situation, it is not difficult to call somatic mutations from paired tumor/normal next generation sequencing data, as only at somatic sites are there bases different from the reference alleles in the tumor genome, but not in the matched normal genome. However, biological and technological factors, including intra-tumor heterogeneity, sample contamination, uncertainties in base sequencing and read alignment, pose a big challenge to somatic mutation discovery [13-15]. Specifically, studies on tumor clonal and sub-clonal structures revealed that tumor cells vary in the way they are abnormal, and some mutations may be observed in only a small fraction of tumor cells in a patient [16, 17]. Furthermore, it is very hard to obtain absolutely pure tumor and normal samples by current experimental technologies, which may result in underestimated variant allele fractions (VAF) in tumor or overestimated VAFs in normal. In addition, technological limitations bring uncertainties in base calling and read alignment. These uncertainties complicate the transformation from aligned data to allelic counts.

A collection of callers and ensembles emerged to detect somatic small mutations from matched tumor/normal, or unmatched tumor sequencing data [18-23]. Designed for the same purpose, callers and ensembles are different in the diversity level of noises taken into account, in the way noises are modelled, in the threshold used to report a mutation as well as in the stringency level to define a false positive in post-call filtering. Validated somatic mutations are valuable resources to evaluate the performance of these callers and guide their improvement. However, it is resource intensive and time consuming to generate ground truth somatic sites [24, 25]. As different sequencing platforms have their own error patterns, multi-platform data from the same sample are needed to complement each other. Standard 30x-50x depths for whole genomes and 100x-150x depths for exomes are not adequate for detection of somatic events in tumors consisting of genetically heterogeneous tumor cells. Deep sequencing is required to offer the desired sensitivity to sub-clonal events. Arbitration is essential for sites whose genotypes disagree between callers or datasets. For the currently available small-sized validated events of individual tumors, they may suffer bias towards one particular validation technology.

Fortunately, simulation of genomic data enables us to generate *in silico* tumors with completely known somatic mutations. Compared with wet-lab validation, computer simulation is much more flexible. Simulated mutations can happen at any genomic site, with any VAF, in any genomic context, and have no limitation in their mutation spectrum. Such flexibilities facilitate characterization of somatic mutation callers and interrogation of their weaknesses. BAMSurgeon is a tool to simulate tumor genomes from normal ones, which was developed by ICGC-TCGA DREAM Somatic Mutation Calling Challenge [26]. It randomly modifies reads spanning the desired sites in the normal or pre-tumor BAM files based on the specified VAFs, and then realigns the modified reads before merging them back into the original BAMs. The modified BAMs serve as tumors with successful spike-in mutations. This kind of simulated tumors are more realistic compared with those from a reference genome assembly, as the underlying biases and error profiles resulting from sequencing technologies and library construction methods are maintained [27]. HapMap/1000 Genomes CEU female NA12878 is the first well-characterized whole-genome reference material from the National Institute of Standards and Technology (NIST) [24, 28]. Its high-confidence genotype calls that include single nucleotide polymorphisms (SNPs), small (1-50 bp) insertions and deletions (indels) and homozygous reference sites have been developed and are publicly available. Furthermore, a large set of sequencing data of NA12878 are freely accessible to researchers [29]. All these lay a foundation for the work presented here.

Our work generated a database of 135 simulated tumor genomes for public use. These simulated tumors were created by BAMSurgeon that introduced small variants (single nucleotide variants (SNVs and small indels) into homozygous reference sites of high confidence of the well-characterized NA12878 genome. To increase the diversity level of sequencing and subsequent mapping errors, we used the NA12878 data (four whole genomes and one exome) from three sequencing centers with different library designs and sample preparations as pre-tumor or normal. Starting with each pre-tumor, 27 increasingly challenging tumors were simulated. The data complexity is displayed in the mutation frequency across the genome, the number of sub-clones and the VAFs. Since local copy number variation (CNV) and tumor ploidy can be computationally generalized to the factor of VAF at each genomic site, we did not include them as factors of data complexity. These pure tumor/normal pairs can be mixed at desired ratios within each pair to further simulate sample contamination. Together, this database of diverse simulated tumors is of great use to benchmark somatic small variant callers’ performance and improve their accuracy metrics.

## Construction and content

### Data sets

We show the procedure about how to develop simulated tumors from pretumors/normals in Figure 1. To enhance analysis of different sequencing and subsequent mapping error profiles, we used four whole genomes and one exome Illumina data for NIST reference material NA12878 as pretumor. Table 1 and Supplemental Material give detailed information about these five pre-tumor/normal sequencing data. Two whole genomes are part of a deep depth (∼300x) dataset of 2x148 paired end reads (which is available at ftp://ftp-trace.ncbi.nlm.nih.gov/giab/ftp/data/NA12878/NIST_NA12878_HG001_HiSeq_300x/) [29]. This dataset was made from 14 libraries in total, and two whole genomes in our work each contained 4 libraries. The other two whole genomes are from a high depth (more than 200x) dataset of 2x100 paired end reads (http://www.ebi.ac.uk/ena/data/view/ERS179577) [28]. The exome data is of 2x100 paired ends and accessible from ftp://ftp-trace.ncbi.nlm.nih.gov/giab/ftp/data/NA12878/Garvan_NA12878_HG001_HiSeq_Exome/. We did not directly use the BAM files provided. Instead, we downloaded the FASTQ format files first, used BWA to map the reads to the human reference genome hg38/GRCh38 with default settings [30], marked PCR and optical duplicates with Picard, realigned the raw gapped alignment and adjusted base quality scores with GATK [31]. The resulting pre-tumor BAM files are NA12878_HiSeq1_normal.bam, NA12878_HiSeq2_normal.bam, NA12878_Exome_normal.bam, NA12878_Illumina1_normal.bam and NA12878_Illumina2_normal.bam. The high-confidence genotype calls for NA12878 are contained in two files: the BED format file that includes the genomic regions whose genotypes were identified confidently and the VCF file for small variants. The genomic sites that are in the BED file and not in the VCF file are homozygous reference allele sites. Such sites have a possibility to get mutated to harbor somatic mutations in the simulated tumors by BAMSurgeon. Currently, two resources of high-confidence genotype calls for NA12878 are available: NIST Genome in a Bottle (GIAB) and Illumina’s Platinum Genomes (PG) [24, 28]. We used the latest GIAB version v3.3.2 call set under GRCh38 available at ftp://ftp-trace.ncbi.nlm.nih.gov/giab/ftp/release/NA12878_HG001/NISTv3.3.2/GRCh38/. For the PG call set, we used the v2016-1.0 under hg38/GRCh38 at ftp://ussd-ftp.illumina.com/2016-1.0/hg38/. We also note the availability of PG v2017-1.0 call set, however, it was not released at the time of preparing our work.

**Figure 1.**
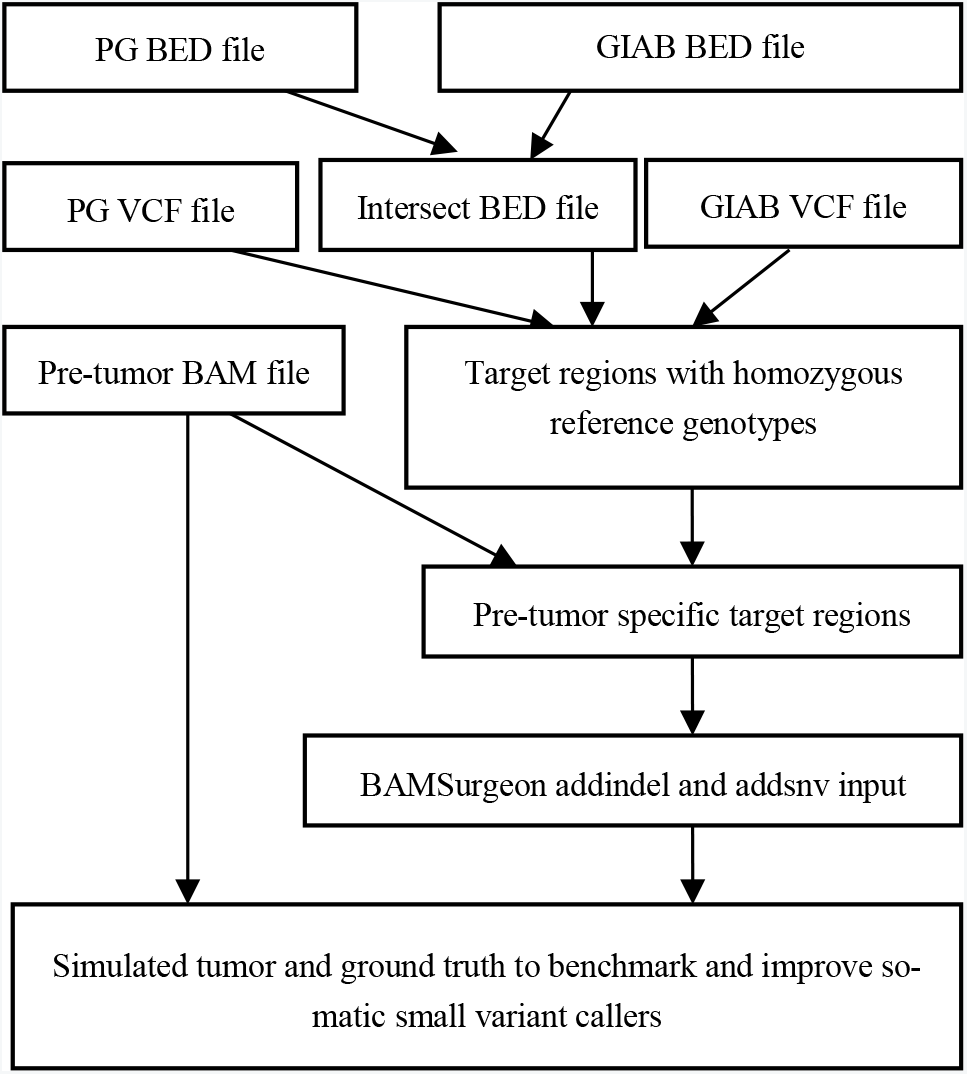
The procedure about how to create simulated tumors by BAM-Surgeon from pre-tumor/normal BAM files to benchmark and tune callers of somatic small variants. It started with PG and GIAB high-confidence genotype calls of NIST reference material NA12878. To further eliminate possible biases towards any particular sequencing technology, read map-per and variant caller used for identifying NA12878 genotype calls, we took only the genomic sites that are shared by both GIAB and PG BED files, and the resulting file is called the intersect BED file. We obtained the target regions with homozygous reference genotypes by masking out the genomic sites with small variants and SNVs in the intersect BED file. Next, for each pre-tumor/normal BAM file, BAMSurgeon randomly selected sites from its target regions for SNV and small indel spike-in. These selected genomic sites in BED format and pre-tumor specific BAM files were provided to BAMSurgeon to yield simulated tumors and ‘truth’ VCF files with successfully spike-in somatic sites. To benchmark and tune somatic small variant callers, a pair of synthetic tumor and its pre-tumor/normal acts as input to return a VCF file with predicted somatic sites, and accuracy metrics are obtained by evaluating the predicted VCF file against its corresponding ground truth VCF file and the benchmark BED file.

**Table 1.**
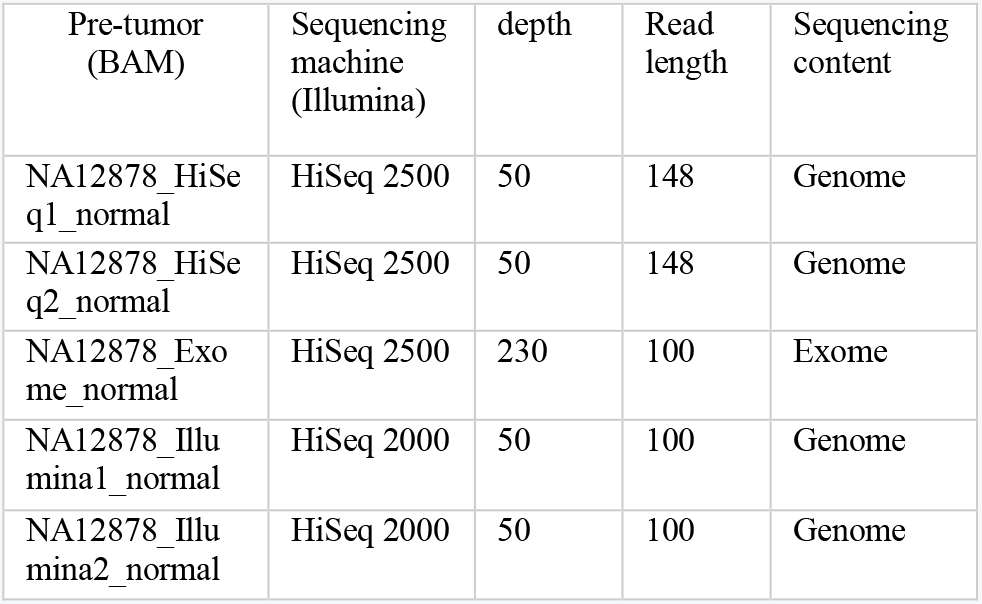
Description of pre-tumor/normal sequencing data of NIST reference material of NA12878 for our simulation work. The pre-tumors/normals are showed by their corresponding BAM file names without extension

### Generating target regions to receive spike-ins

Given two sources of the BED format file consisting of genomic regions with highly confident genotype calls, we used the simple consensus approach to further exclude possible uncertainties, that is, we chose only the genomic sites that are both an element of GIAB and PG BED file. The result is referred to as the intersect BED file here. We then transformed the VCF file from GIAB and PG into the BED format. This step generated the BED file called VCF2BED for each source. The genomic regions that are present only in the intersect BED file and not in the GIAB and PG VCF2BED files have homozygous reference genotypes. They are target regions from which we can randomly select to receive spike-in somatic mutations.

The aforementioned file with target regions was generated without considering the specific sample BAM files. It needs to be modified when it comes to the different pre-tumor sequencing data. For each pre-tumor BAM file, we first calculated the depth of each genomic site within the target regions by SAMtools depth (v1.3.1) [32]. The target regions were filtered by excluding the genomic sites with the depth of lower than 10. The remaining regions were merged by BEDtools merge (v2.26.0) to combine book-end sites or intervals [33], which are the final targets to be played with in the simulation step. The pre-tumor BAM files in the same order as detailed in the subsection of Data sets contain 2295167652, 2294894981, 86922743, 2295481015 and 2296634693 target sites respectively.

### Selecting from target regions for BAMSurgeon spike-in

From each file with final target regions, we used the python script ran-domsites.py in the BAMSurgeon distribution to randomly select genomic sites for SNV spike-in. Then BEDtools subtract (v2.26.0) was performed to extract the genomic sites that are within the final target regions and not selected for SNV spike-ins. The genomic regions from the subtract step were fed into the randomsites.py script for BAMSurgeon addindel input this time. For the input to BAMSurgeon, the randomsites.py generates a list of genomic sites with four columns for addsnv and two more columns for addindel. We filled the fourth column (1-based) with our specified values and kept the other columns untouched. We also kept the default mutated bases returned by BAMSurgeon.

For each pre-tumor, we simulated three types of mutation loads across the genome for SNV spike-ins: 2 mutations per MB (2/MB), 5 mutations per MB (5/MB) and 10 mutations per MB (10/MB). The mutation load for small indel spike-ins was 10% of that of SNVs. There were three types of cell sub-population composition to each mutation load: 2 sub-populations (subclone_2, expected VAFs of 0.5 and 0.35), 3 sub-populations (sub-clone_3, expected VAFs of 0.5, 0.35 and 0.2) and 4 sub-populations (sub-clone_4, expected VAFs of 0.5, 0.35, 0.2 and 0.1). Within a tumor, each sub-population took the same weight. For instance, for a simulated tumor with four sub-populations, four types of mutations in terms of the VAF each represented 25% of the total number. To create mutations in a diversity of genomics contexts, we performed three random selections for the same characteristics (mutation frequency across the genome and the number of cell sub-populations). These parameters gave 27 simulated tumors in total from each pre-tumor.

### Simulating somatic small variants with BAMSurgeon

We fed pre-tumor BAM files and their corresponding addsnv input to BAMSurgeon, and this step created simulated tumors with somatic SNVs. The simulated tumors from this step have some records that do not respect the sorting order of BAM file. So we resorted by position and indexed the BAM files by SAMtools (v1.3.1). Then they were fed into BAMSurgeon as the value of option -f with their corresponding addindel input to receive somatic small indels. The resulting BAM files were resorted and indexed before they could be used for benchmarking somatic small variant callers. The parameters used when running BAMSurgeon to generate addsnv input, addindel input and simulated tumors are provided in Supplemental Material.

Take the simulated tumor BAM file NA12878_1_snv_indel_sorted.bam for instance, which has 2 sub-clones (expected VAFs of 0.5 and 0.35). Its somatic SNV and small indel mutation frequency are 2/MB and 0.2/MB respectively. To generate this tumor, we used BAMSurgeon to modify the pre-tumor/normal BAM file NA12878_HiSeq1_normal.bam based on the output of the randomsites.py script for BAMSurgeon addsnv input. The output was NA12878_1_snv.bam and snv_1.vcf. The BAM file was resorted and indexed to be NA12878_1_snv_sorted.bam. Somatic small indels in the output of the randomsites.py script for BAMSurgeon addindel input were added to it to yield NA12878_1_snv_indel.bam and indel_1.vcf. The operations of resorting and indexing were performed on this BAM file and made the final tumor NA12878_1_snv_indel_sorted.bam available. The snv_1.vcf and indel_1.vcf from BAMSurgeon were unordered, so we sorted them using natural ordering by vcf-sort (v0.1.15) [34]. Then the sorted VCF files were bgzipped (v1.2. and tabix indexed by SAMtools (v1.3.1) before being provided to vcf-merge (v0.1.15) to get merged by genomic position. The resulting snv_indel_1.vcf file contains ground truth somatic SNVs and small indels in the simulated tumor NA12878_1_snv_indel_sorted.bam. Researchers need both of them and the matched pre-tumor/normal NA12878_HiSeq1_normal.bam for evaluation and benchmark.

### Benchmarking with simulated tumors

To accurately benchmark somatic callers, we need to focus on just the regions with highly confident genotype calls. Besides, to comprehensively benchmark somatic callers in terms of accuracy metrics, every genomic site in the focused regions should have one of the three genotype calls: wildtype/reference site, germline site and somatic site. In the context of benchmarking with simulated tumors, successfully spike-in somatic small variants are positives to calculate sensitivity, and the rest of sites (germlines and wildtypes) are negatives to calculate specificity. For each of the pre-tumors/normals, we generated a benchmark BED file named NA12878_HiSeq1_benchmark.bed, NA12878_HiSeq2_benchmark.bed, NA12878_Exome_benchmark.bed, NA12878_Illumina1_benchmark.bed and NA12878_Illumina2_benchmark.bed. The benchmark BED files were generated by intersecting the aforementioned intersect BED file with each pre-tumor/normal BAM file. To benchmark and tune somatic small variant callers, a pair of simulated tumor and its pre-tumor/normal are inputs to output a VCF file with predicted somatic sites. Sensitivity is calculated by evaluating the predicted VCF file against its corresponding ground truth VCF file, and the remaining sites that are in the benchmark BED file but not in the ground truth VCF file are used to calculate specificity.

## Utility and discussion

### Overview of the database of simulated tumors

Our work is motivated by the lack of ground truth somatic mutations to benchmark somatic callers. It is the product of the characterization of NIST reference material NA12878 genome and state-of-the-art simulation tool BAMSurgeon. Two projects have been working on identifying genotypes of NA12878 with high accuracy: NIST GIAB and Illumina’s PG. Their own version’s genotype calls are publicly available on their corresponding websites. These genotypes calls include high-confidence SNPs, small indels and homozygous reference sites. BAMSurgeon simulates tumors by introducing synthetic mutations to original genomes. It modifies the reads spanning genomic sites to get these sites mutated. The reads covering the desired sites each have a probability that is equal to the user-specified VAF to be selected and get modified into the variant allele. This way, simulated tumors are realistic and keep error profiles from library preparation and sequencing machines.

Our website contains 135 simulated tumors, which were yielded by introducing small variants into the homozygous reference sites or wildtype sites of NA12878 genome by BAMSurgeon. Every 27 of them were created from one normal BAM file. Figure 2 shows the organization of these simulated tumors on our website, and the file Simulated_tumor_information there gives detailed information about the characteristics of these tumors. The fold Ground_truth_VCF_files comprises all the ground truth files with somatic SNVs and small indels. The rest five folds are named after the pre-tumor/normal BAM file without the extension. Within them are the pre-tumor/normal BAM file and 27 corresponding simulated tumors. These tumor files have names from NA12878_1_snv_indel_sorted.bam to NA12878_135_snv_indel_sorted.bam. To benchmark somatic callers, a pre-tumor/normal BAM file, one simulated tumor from 27 files and the corresponding ground truth VCF file with the same index/number as that of the simulated tumor are needed. Within one fold, these tumors are different in the number of sub-clones, the VAF, the mutation frequency across the genome and the mutation site. Between folds, simulated tumors are different in sequencing and subsequent mapping error profiles, read length and capture content.

**Figure 2.**
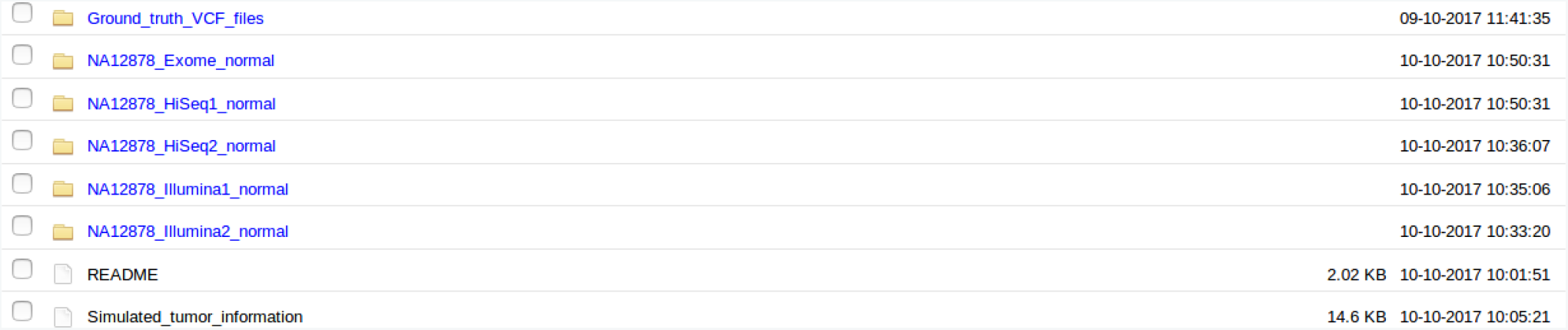
The organization of simulated tumors on our website. The file Simulated_tumor_information provides detailed information on the characteristics of these tumors. The fold Ground_truth_VCF_files comprises all the ground truth files with somatic SNVs and small indels. The remaing five folds are named after the pre-tumor/normal BAM files without the extension, and each of them includes the pre-tumor/normal BAM file, benchmark BED file and 27 corresponding simulated tumors.

### Genomic features of spike-in somatic sites

Due to the parameters’ constrain in BAMSurgeon, not all chosen spikeins can be successful. Supplemental Table S2 provides the successful rate of somatic SNV and small indel spike-ins for each simulated tumor. The overall successful rate for somatic small indel spike-ins is higher than that of somatic SNV spike-ins. We chose 1402598 somatic SNV sites and 141743 somatic small indel sites respectively. 1285692 somatic SNV sites were mutated successfully (successful rate is about 0.917). The successful rate is about 0.991 for somatic small indels. Of the 135 simulated tumors, the minimum, median and maximum successful rate are 0.887, 0.932 and 0.960 for somatic SNV spike-ins, and 0.889, 0.989 and 1 for somatic small indel spike-ins, respectively.

Considering the importance of genomic context in somatic SNV calling, we extracted the three bases (from −1 bp to +1 bp) centered on the simulated somatic sites of hg38 reference genome for successful and unsuccessful somatic SNV spike-ins. Figure 3 shows the fraction of 64 trinucleotide contexts for somatic SNV spike-ins. Successful and unsuccessful somatic SNV sites display similar distributions of trinucleotide contexts. AAA and TTT have the highest proportions for both types of somatic SNV sites. On the contrary, TCG, CGT, CGA and ACG are the least contributors, with a proportion of approximately 0.005.

**Figure 3.**
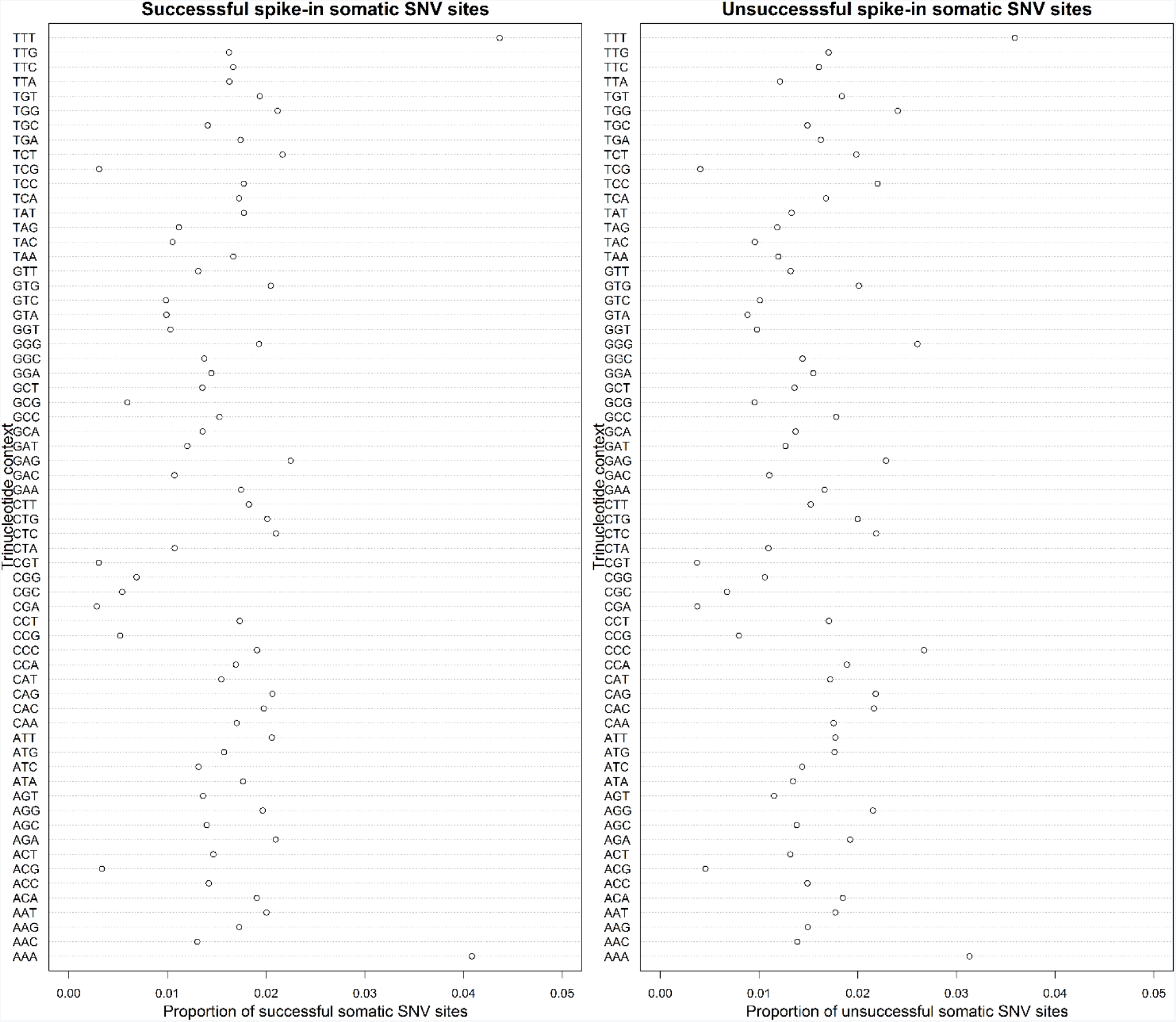
The fraction of 64 trinucleotide contexts for successful somatic SNV sites (left panel) and unsuccessful ones (right panel). Three bases (from −1 bp to +1 bp) centered on the simulated somatic sites of hg38 reference genome were extracted for successful and unsuccessful somatic SNV spike-ins respectively. Then we calculated the proportion by dividing the number of somatic SNVs with a type of trinucleotide context by the total number.

Then to determine the category of known repeats to which the simulated sites belong, we downloaded the RepeatMasker-masked regions rmsk.txt.gz from http://hgdownload.cse.ucsc.edu/goldenpath/hg38/database, which contains a detailed annotation of the repeats present in hg38 human reference genome and was generated by Arian Smit’s Repeat-Masker Program at http://www.repeatmasker.org/. The format description of file rmsk.txt is available at http://genome.ucsc.edu/cgi-bin/hgTables. We mapped the genomic sites of successful and unsuccessful somatic SNVs and small indels to the regions in rmsk.txt by BEDOPS bedmap [35]. Tables 2 and 3 give the detailed composition of 16 different types of repeat sequence and non-repeat sequence (others). For successful somatic sites, approximately half of them occur at regions of non-repeat sequence, 0.4374 for somatic SNV sites and 0.4120 for somatic small indel sites. When it comes to unsuccessful somatic sites, only approximately 0.15 of them lie within regions of non-repeat sequence, and almost half of them (0.5210 and 0.4814 for somatic SNVs and small indels respectively) come from regions of LINE of repeat sequence.

**Table 2.**
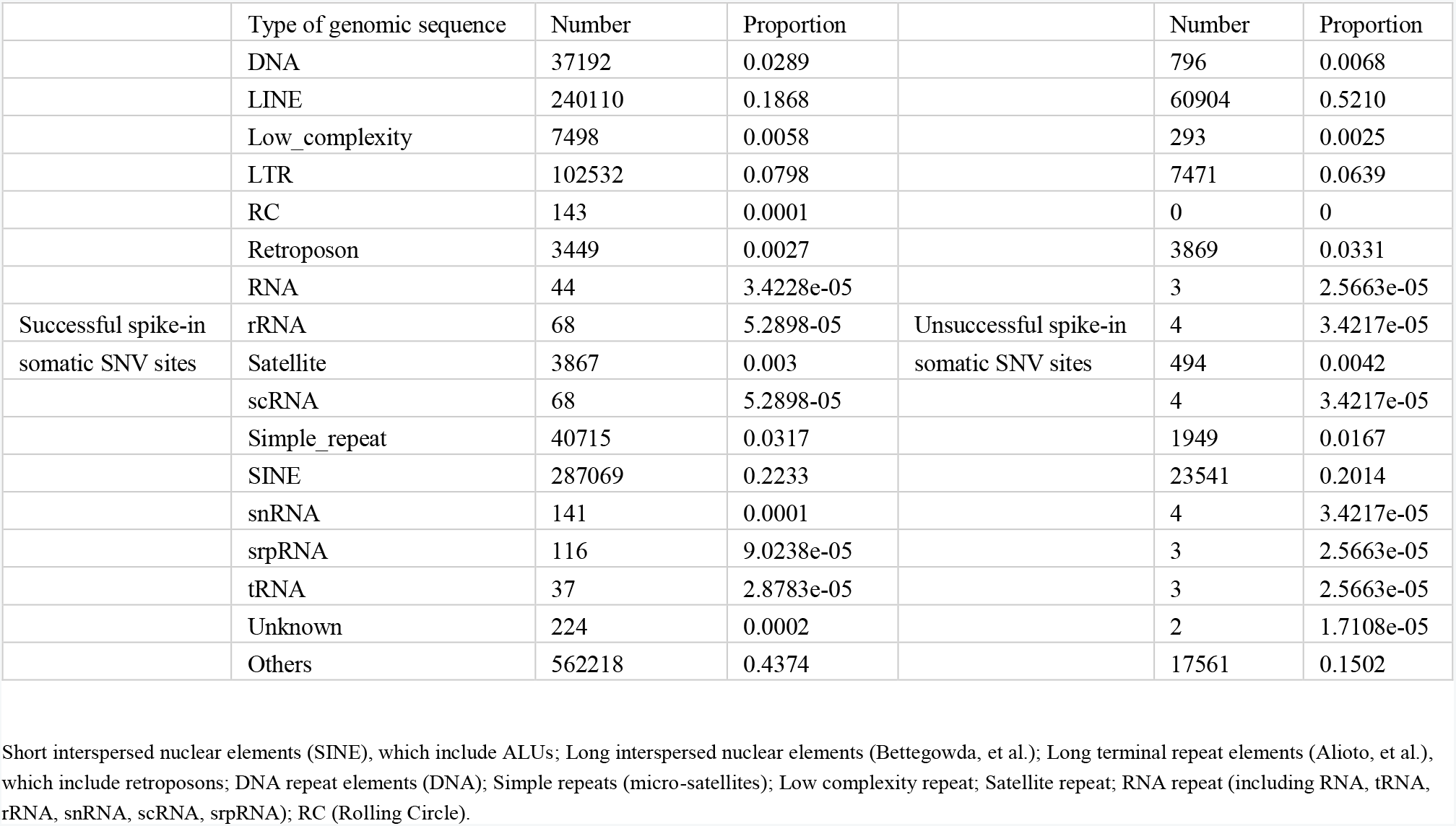
The category of known repeats in RepeatMasker regions and non-repeat sequence to which the simulated somatic SNV sites belong. Others stands for non-repeat sequence.

|  | Type of genomic sequence | Number | Proportion |  | Number | Proportion |
| --- | --- | --- | --- | --- | --- | --- |
|  | DNA | 37192 | 0.0289 |  | 796 | 0.0068 |
|  | LINE | 240110 | 0.1868 |  | 60904 | 0.5210 |
|  | Low_complexity | 7498 | 0.0058 |  | 293 | 0.0025 |
|  | LTR | 102532 | 0.0798 |  | 7471 | 0.0639 |
|  | RC | 143 | 0.0001 |  | 0 | 0 |
|  | Retroposon | 3449 | 0.0027 |  | 3869 | 0.0331 |
|  | RNA | 44 | 3.4228e-05 |  | 3 | 2.5663e-05 |
| Successful spike-in | rRNA | 68 | 5.2898e-05 | Unsuccessful spike-in | 4 | 3.4217e-05 |
| somatic SNV sites | Satellite | 3867 | 0.003 | somatic SNV sites | 494 | 0.0042 |
|  | scRNA | 68 | 5.2898e-05 |  | 4 | 3.4217e-05 |
|  | Simple_repeat | 40715 | 0.0317 |  | 1949 | 0.0167 |
|  | SINE | 287069 | 0.2233 |  | 23541 | 0.2014 |
|  | snRNA | 141 | 0.0001 |  | 4 | 3.4217e-05 |
|  | srpRNA | 116 | 9.0238e-05 |  | 3 | 2.5663e-05 |
|  | tRNA | 37 | 2.8783e-05 |  | 3 | 2.5663e-05 |
|  | Unknown | 224 | 0.0002 |  | 2 | 1.7108e-05 |
|  | Others | 562218 | 0.4374 |  | 17561 | 0.1502 |

Short interspersed nuclear elements (SINE), which include ALUs; Long interspersed nuclear elements (Bettegowda, et al.); Long terminal repeat elements (Alioto, et al.), which include retroposons; DNA repeat elements (DNA); Simple repeats (micro-satellites); Low complexity repeat; Satellite repeat; RNA repeat (including RNA, tRNA, rRNA, snRNA, scRNA, srpRNA); RC (Rolling Circle).

**Table 3.** The category of known repeats in RepeatMasker regions and non-repeat sequence to which the simulated somatic small indel sites belong. Others stands for non-repeat sequence.

|  | Type of genomic sequence | Number | Proportion |  | Number | Proportion |
| --- | --- | --- | --- | --- | --- | --- |
|  | DNA | 3821 | 0.0273 |  | 7 | 0.0052 |
|  | LINE | 29805 | 0.2127 |  | 647 | 0.4814 |
|  | Low_complexity | 817 | 0.0058 |  | 2 | 0.0015 |
|  | LTR | 11039 | 0.0788 |  | 46 | 0.0342 |
|  | RC | 20 | 0.0001 |  | 0 | 0 |
|  | Retroposon | 732 | 0.0052 |  | 31 | 0.0231 |
|  | RNA | 8 | 5.7103e-05 |  | 0 | 0 |
| Successful spike-in | rRNA | 3 | 2.1414e-05 | Unsuccessful spike-in | 0 | 0 |
| somatic small indel sites | Satellite | 453 | 0.0032 | somatic small indel sites | 1 | 0.0007 |
|  | scRNA | 11 | 7.8517e-05 |  | 0 | 0 |
|  | Simple_repeat | 4492 | 0.0321 |  | 7 | 0.0052 |
|  | SINE | 31119 | 0.2221 |  | 394 | 0.2932 |
|  | snRNA | 19 | 0.0001 |  | 0 | 0 |
|  | srpRNA | 11 | 7.8517e-05 |  | 0 | 0 |
|  | tRNA | 2 | 1.4276e-05 |  | 0 | 0 |
|  | Unknown | 22 | 0.0002 |  | 0 | 0 |
|  | Others | 57723 | 0.4120 |  | 209 | 0.1555 |

Short interspersed nuclear elements (SINE), which include ALUs; Long interspersed nuclear elements (Bettegowda, et al.); Long terminal repeat elements (Alioto, et al.), which include retroposons; DNA repeat elements (DNA); Simple repeats (micro-satellites); Low complexity repeat; Satellite repeat; RNA repeat (including RNA, tRNA, rRNA, snRNA, scRNA, srpRNA); RC (Rolling Circle).

### Benchmark BED files

Due to the limitations of current sequencing technologies, read mappers and variant callers, it is impossible to identify the genotype call of every genomic site across the whole genome. Thus, to accurately and comprehensively benchmark somatic small variant callers, we need to focus on just the regions where every genomic site has one of the three high-confidence genotypes: wildtype or reference site, germline site and somatic site. To meet the purpose, we created a benchmark BED file for each of the pre-tumors/normals, named NA12878_HiSeq1_benchmark.bed, NA12878_HiSeq2_benchmark.bed, NA12878_Exome_benchmark.bed, NA12878_Illumina1_benchmark.bed and NA12878_Illumina2_bench mark.bed, which contain 2302950972, 2302925972, 991374103, 2303071124 and 2303135374 genomic sites respectively. When doing benchmark, somatic small variant callers take in a pair of simulated tumor and its pre-tumor/normal, and yield a VCF file with predicted somatic sites. The VCF file and its corresponding ground truth VCF file are used to calculate sensitivity. The negatives are the remaining genomic sites that are in the benchmark BED file but not in the ground truth VCF file, which are used to calculate specificity.

## Conclusions

Accurate detection of somatic sites is critical to clinical diagnosis. A lot of somatic callers have been developed so far to identify somatic small variants from matched tumor/normal, or unmatched tumor sequencing data. However, the limited number of validated somatic sites challenges the evaluation and then improvement of somatic callers. Fortunately, computing simulation of genomic data makes it possible to create simulated tumors with ground truth somatic mutations.

Genotype calls with high confidence of NIST reference material NA12878 genome are publicly available. The genotype calls consist of SNPs, small indels and homozygous reference sites. Also, different types of sequencing data of NA12878 are freely accessible to researchers. Given these available resources corresponding to NA12878, our work introduced somatic variants into homozygous reference or wildtype sites of its genome. BAMSurgeon performed the work of introducing somatic variants by modifying the reads covering the chosen genomic sites. This way, simulated tumors are more realistic, as they maintain the underlying error profiles stemming from library construction methods, sequencing technologies and then mapping algorithms.

We created 135 simulated tumors with somatic SNVs and small indels in total from 5 pre-tumor/normal BAM files. These pre-tumors/normals are different from each other in sequencing and subsequent mapping error profiles, read length, the number of sub-clones, the VAF, the mutation frequency across the genome and the genomic feature. Furthermore, to evaluate somatic callers’ performance in the situation of sample contamination, contaminated samples can be simulated by mixing these pure tumor/normal pairs at desired ratios. Together, this database of simulated tumors with high diversity will be a valuable resource to benchmark somatic small variant callers and guide their improvement.

## Declarations

### Ethics approval and consent to participate

Not applicable.

### Consent for publication

Not applicable.

### Availability of data and material

The database is freely available at ftp://137.92.56.98/simulated-tumors and https://trace.ncbi.nlm.nih.gov/Traces/sra/sra.cgi?study=SRP115159.

## Acknowledgements

We would like to thank National Computational Infrastructure of Australia for its High Performance Computing (HPC) Systems and Cloud Computing Systems.

